# anndata: Annotated data

**DOI:** 10.1101/2021.12.16.473007

**Authors:** Isaac Virshup, Sergei Rybakov, Fabian J. Theis, Philipp Angerer, F. Alexander Wolf

## Abstract

anndata is a Python package for handling annotated data matrices in memory and on disk (github.com/theislab/anndata), positioned between pandas and xarray. anndata offers a broad range of computationally efficient features including, among others, sparse data support, lazy operations, and a PyTorch interface.

**Statement of need:** Generating insight from high-dimensional data matrices typically works through training models that annotate observations and variables via low-dimensional representations. In exploratory data analysis, this involves *iterative* training and analysis using original and learned annotations and task-associated representations. anndata offers a canonical data structure for book-keeping these, which is neither addressed by pandas (McKinney, 2010), nor xarray (Hoyer & Hamman, 2017), nor commonly-used modeling packages like scikit-learn (Pedregosa et al., 2011).

## Introduction

Since its initial publication as part of Scanpy (Wolf et al., 2018), anndata matured into an independent software project and became widely adopted (694k total PyPI downloads & 48k downloads/month, 225 GitHub stars & 581 dependent repositories).

anndata has been particularly useful for data analysis in computational biology where advances in single-cell RNA sequencing (scRNA-seq) gave rise to new classes of analysis problems with a stronger adoption of Python over the traditional R ecosystem. Previous bulk RNA datasets had few observations with dense measurements while more recent scRNA-seq datasets come with high numbers of observations and sparse measurements, both in 20k dimensions and more. These new data profit much from the application of the scalable machine learning tools of the Python ecosystem.

## The AnnData object

`AnnData` is designed for data scientists and was inspired by a similar data structure in the R ecosystem, `ExpressionSet` (Huber et al., 2015).

Within the pydata ecosystem, xarray (Hoyer & Hamman, 2017) enables to deal with labeled data tensors of arbitrary dimensions, while pandas (McKinney, 2010) operates on single data matrices (tables) represented as `DataFrame` objects. anndata is positioned in between pandas and xarray by providing structure that organizes data matrix annotations. In contrast to pandas and xarray, `AnnData` offers a native on-disk format that allows sharing data with analysis results in form of learned annotations.

### The data structure

Standardized data structures facilitate data science, with one of the most adopted standards being *tidy data* (Wickham, 2014). anndata complies with *tidy data* but introduces additional conventions by defining a data structure that makes use of conserved dimensions between data matrix and annotations. With that, `AnnData` makes a particular choice for data organization that has been left unaddressed by packages like scikit-learn or PyTorch (Paszke et al., 2019), which model input and output of model transformations as unstructured sets of tensors.

At the core of `AnnData` is the measured data matrix from which we wish to generate insight (`X`). Each data matrix element stores a value and belongs to an observation in a row (`obs_names`) and a variable in a column (`var_names`), following the *tidy data* standard. Performing exploratory data analysis with `AnnData`, one builds an understanding of the data matrix by annotating observations and variables using `AnnData`’s fields (Figure 1) as follows:

**Figure 1:**
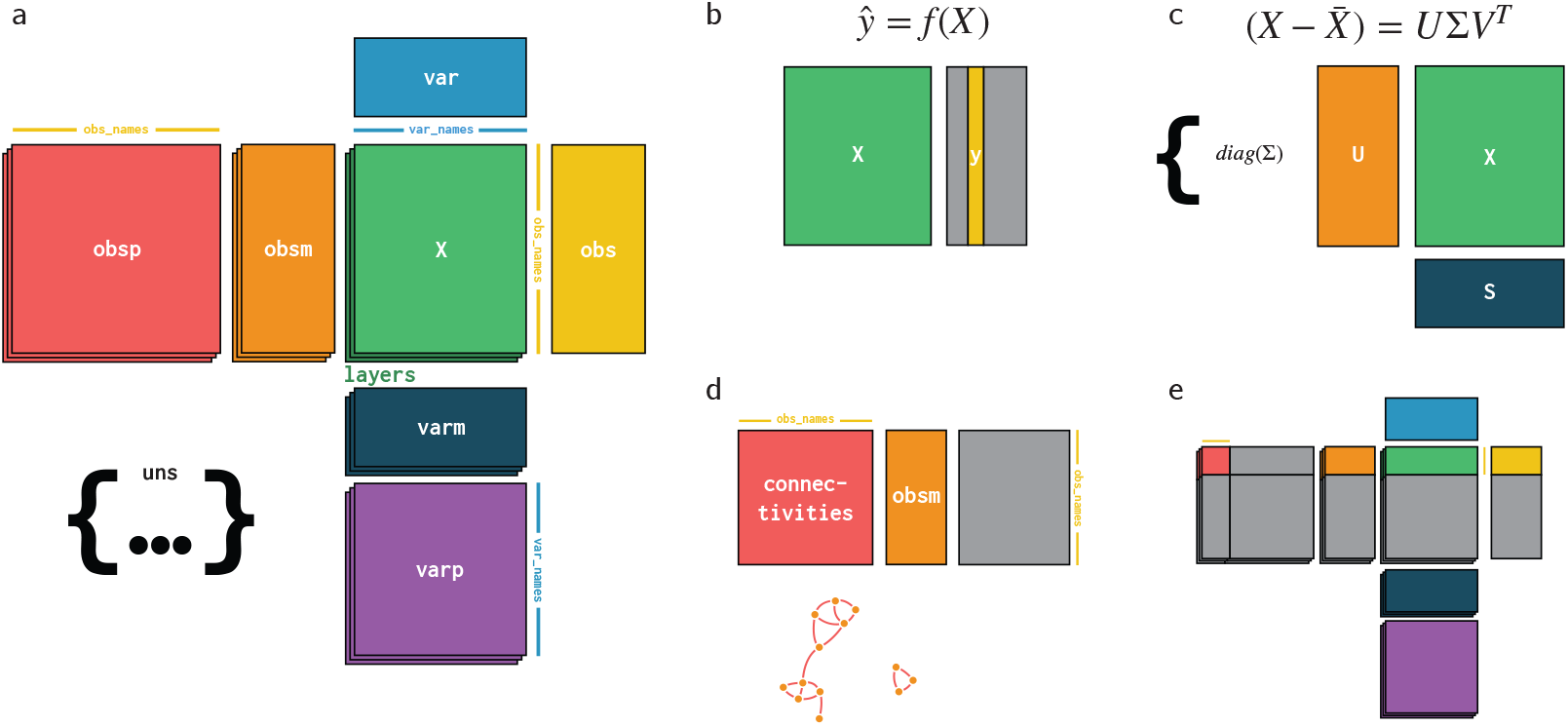
Structure of the AnnData object. **a**, The AnnData object is a collection of arrays aligned to the common dimensions of observations (`obs`) and variables (`var`). Here, color is used to denote elements of the object, with “warm” colors selected for elements aligned to the observations and “cool” colors for elements aligned to variables. The object is centered around the main data matrix `X`, whose two dimensions correspond to observations and variables respectively. Primary labels for each of these dimensions are stored as `obs_names` and `var_names`. If needed, `layers` stores matrices of the exact same shape as `X`. One-dimensional annotations for each dimension are stored in pandas `DataFrame`s `obs` and `var`. Multi-dimensional annotations are stored in `obsm` and `varm`. Pairwise relationships are stored in `obsp` and `varp`. Unstructured data which doesn’t fit this model, but should stay associated to the dataset are stored in `uns`. **b**, Let us discuss a few examples. The response variable ŷ learned from X is stored as a one-dimensional annotation of observations. **c**, Principal components and the transformed dimensionality-reduced data matrix obtained through PCA can be stored as multi-dimensional annotations of variables and observations, respectively. **d**, A k-nearest neighbor graph of any desired representation is stored as a sparse adjacency matrix of pairwise relationships among observations in `obsp`. This is useful to have easy access to the connectivities of points on a low-dimensional manifold. **e**, Subsetting the `AnnData` object by observations produces a view of data and annotations.

- One-dimensional annotations get added to the main annotation `DataFrame` for each axis, `obs` and `var`.
- Multi-dimensional representations get added to `obsm` and `varm`.
- Pair-wise relations among observations and variables get added to `obsp` and `varp` in form of sparse graph adjacency matrices.

Prior annotations of observations will often denote the experimental groups and conditions that come along with measured data. Derived annotations of observations might be summary statistics, cluster assignments, low-dimensional representations or manifolds. Annotations of variables will often denote alternative names or measures quantifying feature importance.

In the context of how (Wickham, 2014) recommends to order variables, one can think of `X` as contiguously grouping the data of a specific set of *measured* variables of interest, typically high-dimensional readout data in an experiment. Other tables aligned to the observations axis in `AnnData` are then available to store both *fixed* (meta-)data of the experiment and derived data.

We note that adoption of *tidy data* (Wickham, 2014) leaves some room for ambiguity. For instance, the R package `tidySummarizedExperiment` (Mangiola, 2021) provisions tables for scRNA-seq data that take a long form that spreads variables belonging to the same observational unit (a cell) across multiple rows. Generally, it may occur that there is no unique observational unit that is defined through a *joint measurement*, for instance, by measuring variables in the same system at the same time. It such cases, the *tidy data* layout is ambiguous and results in longer or wider table layouts depending on what an analyst considers the observational unit.

### The data analysis workflow

Let us illustrate how `AnnData` supports analysis workflows of iteratively learning representations and scalar annotations. For instance, training a clustering, classification or regression model on raw data in `X` produces an estimate of a response variable *ŷ*. This derived vector is conveniently kept track off by adding it as an annotation of observations (`obs`, Figure 1b). A reduced dimensional representation obtained through, say Principal Component Analysis or any bottleneck layer of a machine learning model, would be stored as multi-dimensional annotation (`obsm`, Figure 1c). Storing low-dimensional manifold structure within a desired reduced representation is achieved through a k-nearest neighbor graph in form of a sparse adjacency matrix: a matrix of pairwise relationships of observations (`obsp`, Figure 1d). Subsetting the data by observations produces a memory-efficient view of `AnnData` (Figure 1e).

### The efficiency of data operations

Due to the increasing scale of data, we emphasized efficient operations with low memory and runtime overhead. To this end, anndata offers sparse data support, out of core conversions between dense and sparse data, lazy subsetting (“views”), per-element operations for low total memory usage, in-place subsetting, combining `AnnData` objects with various merge strategies, lazy concatenation, batching, and a backed out-of-memory mode.

In particular, `AnnData` takes great pains to support efficient operations with sparse data. While there is no production-ready API for working with sparse and dense data in the python ecosystem, `AnnData` abstracts over the existing APIs making it much easier for novices to handle each. This concerns handling data both on-disk and in-memory with operations for out-of-core access. When access patterns are expected to be observation/row-based as in batched learning algorithms, the user can store data matrices as CSR sparse matrices or C-order dense matrices. For access along variables, for instance, to visualize gene expression across a dataset, CSC sparse and Fortran order dense matrices allow fast access along columns.

### The on-disk format

An `AnnData` object captures a unit of the data analysis workflow that groups original and derived data together. Providing a persistent and standard on-disk format for this unit relieves the pain of working with many competing formats for each individual element and thereby aids reproducibility. This is particularly needed as even pandas `DataFrame` has no canonical persistent data storage format. `AnnData` has chosen the self-describing hierarchical data formats HDF5 (Collette, 2013) and zarr (Miles et al., 2020) for this purpose (Figure 2), which are compatible with non-Python programming environments. The broad compatibility and high stability of the format led to wide adoption, and initiatives like the Human Cell Atlas (Regev et al., 2017) and HuBMAP (Consortium & others, 2019) distribute their single-cell omics datasets through `.h5ad`.

**Figure 2:**
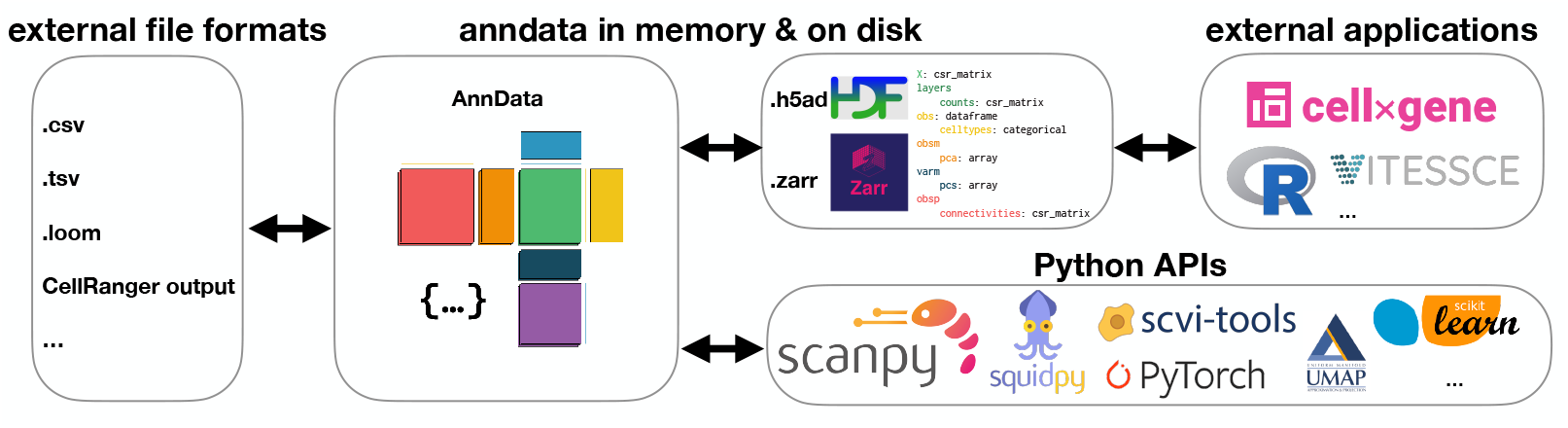
AnnData provides broad interoperability with tools and platforms. `AnnData` objects can be created from a number of formats, including common delimited text files, or domain-specific formats like `loom` files or `CellRanger` outputs. Once in memory, AnnData provides an API for handling annotated matrices, proving a common base object used by a range of analytic computational biology tools and integrating well with the APIs of the established Python machine learning ecosystem. The in memory format has a one-to-one relationship with its hierarchical on disk formats (mapping of elements indicated by color) and uses language-independent technologies, enabling the use by non-Python applications and interchange with other ecosystems.

Within HDF5 and zarr, we could not find a standard for sparse matrices and `DataFrame` objects. To account for this, we defined a schema for these types, which specifies how these elements can be read from disk to memory. This schema is versioned and stored in an internal registry, which evolves with anndata while maintaining the ability to access older versions. On-disk formats within this schema closely mirror their in-memory representations: Compressed sparse matrices (CSR, CSC) are stored as a collection of three arrays, `data`, `indices`, and `indptr`, while tabular data is stored in a columnar format.

## The ecosystem

Over the past 5 years, an ecosystem of packages that are built around anndata has grown. This ecosystem is highly focused on scRNA-seq (Figure 2), and ranges from Python APIs (Zappia & Theis, 2021) to user-interface-based applications (Megill et al., 2021). Tools like scikit-learn and UMAP (McInnes et al., 2020), which are designed around numpy and not anndata, are still centered around data matrices and hence integrate seamlessly with anndata-based workflows. Since releasing the PyTorch `DataLoader` interface `AnnLoader` and the lazy concatenation structure `AnnCollection`, `anndata` also offers native ways of integrating into the Pytorch ecosystem. scvi-tools (Gayoso et al., 2021) offers a widely used alternative for this.

Through the language-independent on-disk format `h5ad`, interchange of data with non-Python ecosystems is easily possible. For analysis of scRNA-seq data in R this has been further simplified by anndata2ri, which allows conversion to `SingleCellExperiment` (Amezquita et al., 2020) and Seurat’s data format (Hao et al., 2020).

Let us give three examples of `AnnData`’s applications: spatial transcriptomics, multiple modalities, and RNA velocity (Figure 3). In spatial transcriptomics, each high-dimensional observation is annotated with spatial coordinates. Squidpy (Palla et al., 2021) uses `AnnData` to model their data by storing spatial coordinates as an array (`obsm`) and a spatial neighborhood graph (`obsp`), which is used to find features which are spatially correlated (Figure 3a). In addition, values from the high-dimensional transcriptomic measurement can be overlaid on an image of the sampled tissue, where an image array (reference) is stored in `uns`.

**Figure 3:**
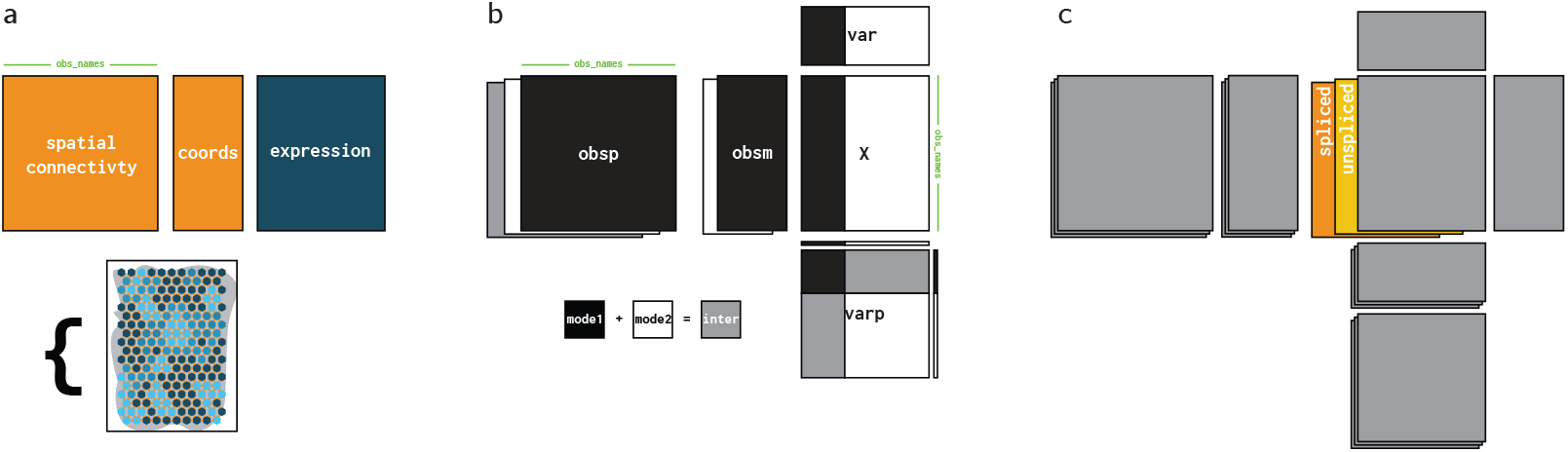
Examples of how AnnData is used by packages in the ecosystem. **a**, Squidpy uses `AnnData` for working with spatial data: the coordinates of each observation are stored in `obsm`, a spatial neighborhood graph in `obsp`, and a complementary image is stored in `uns`. **b**, Multiple modalities can be represented in multiple `AnnData` objects. The variables axis now corresponds to the union of variables across modalities. Modality-specific and joint representations and manifolds are stored as separate elements in `obsm` or `obsp`, while inter-modality relations can be stored as graphs in `varp`. **c**, `AnnData` allows for RNA velocity analyses by storing counts of different splicing states as separate `layers` with velocity-based directed graphs in `obsp`.

To model multimodal data, one approach is to join separate `AnnData` objects (Figure 3b) for each modality on the observations index through `anndata.concat`. Relations between the variables of different modalities can then be stored as graphs in `varp`, and analyses using information from both modalities, like a joint manifold, in `obsp`. Formalizing this further, the `muon` package (Bredikhin et al., 2021) offers a container-like object `MuData` for a collection of `AnnData` objects, one for each modality. This structure extends to an on-disk format where individual `AnnData` objects are stored as discrete elements inside `h5mu` files. This approach has similarity with `MultiAssayExperiment` within the Bioconductor ecosystem (Ramos et al., 2017).

`AnnData` has been used to model data for fitting models of RNA velocity (Bergen et al., 2020) exploiting the `layers` field to store a set of matrices for different types of RNA counts (Figure 3c).

## Outlook

The anndata project is under active development towards more advanced out-of-core access, better cloud & relational database integration, a split-apply-combine framework, and interchange with more formats, like Apache Arrow or TileDB (Papadopoulos et al., 2016). Furthermore, anndata engages with projects that aim at building out infrastructure for modeling multi-modal (Bredikhin et al., 2021) and non-homogeneous data, for instance, to enable learning from Electronic Health Records (Heumos & Theis, 2021). Finally, we aim at further complementing anndata by interfacing with scientific domain knowledge and data provenance tracking.

## Acknowledgements

Isaac is grateful to Christine Wells for consistent support and freedom to pursue work on anndata and Scanpy. We thank Ryan Williams and Tom White for contributing code related to zarr and Jonathan Bloom for contributing a comprehensive PR on group-by functionality. Alex and Phil thank Cellarity for supporting continued engagement with open source software. We are grateful to Fabian’s lab for continuing dissemination along with Scanpy over the past years. This project receives funding through CZI’s Essential Open Source Software for Science grant.

## Author contributions

Isaac has led the anndata project since v0.7, and contributed as a developer before. His contributions include generalized storage for sparse matrices, IO efficiency, dedicated graph storage, concatenation, and general maintenance. Sergei made diverse contributions to the code base, in particular, the first version of `layers`, benchmarking and improvement of the earlier versions of the IO code, the PyTorch dataloader `AnnLoader` and the lazy concatenation data structure `AnnCollection`. Fabian contributed to supervision of the project. Phil co-created the package. He suggested to replace Scanpy’s initial unstructured annotated data object to one mimicking R’s ExpressionSet, and wrote AnnData’s [first implementation](https://github.com/theislab/scanpy/commit/315859c5586116434ea3b7ce97512a5e2a1030e2) with indexing and slicing affecting one-dimensional metadata and the central matrix. He further ascertained good software practices in the project, authored the documentation tool extensions for scanpy and anndata and anndata2ri, a library for in-memory conversion between anndata and SingleCellExperiment. Alex co-created the package. He introduced centering data science workflows around an [initially unstructured annotated data object](https://github.com/theislab/scanpy/tree/c22e48abe45a6ccca5918bbf689637caa4b31250), designed the API, wrote tutorials and documentation until v0.7, and implemented most of the early functionality, among others, reading & writing, the on-disk format `h5ad`, views, sparse data support, concatenation, backed mode. Isaac and Alex wrote the paper with help from all co-authors.

## References

Amezquita, R. A., Lun, A. T., Becht, E., Carey, V. J., Carpp, L. N., Geistlinger, L., Marini, F., Rue-Albrecht, K., Risso, D., Soneson, C., & others. (2020). Orchestrating single-cell analysis with bioconductor. Nature Methods, 17 (2), 137–145.

Bergen, V., Lange, M., Peidli, S., Wolf, F., & Theis, F. (2020). Generalizing RNA velocity to transient cell states through dynamical modeling. Nature Biotechnology, 38(12), 1408–1414.

Bredikhin, D., Kats, I., & Stegle, O. (2021). Muon: Multimodal omics analysis framework. bioRxiv. https://doi.org/10.1101/2021.06.01.445670

Collette, A. (2013). Python and HDF5. O’Reilly.

Consortium, H., & others. (2019). The human body at cellular resolution: The NIH human biomolecular atlas program. Nature, 574(7777), 187.

Gayoso, A., Lopez, R., Xing, G., Boyeau, P., Wu, K., Jayasuriya, M., Melhman, E., Langevin, M., Liu, Y., Samaran, J., Misrachi, G., Nazaret, A., Clivio, O., Xu, C., Ashuach, T., Lotfollahi, M., Svensson, V., Beltrame, E. da V., Talavera-López, C., … Yosef, N. (2021). Scvi-tools: A library for deep probabilistic analysis of single-cell omics data. bioRxiv. https://doi.org/10.1101/2021.04.28.441833

Hao, Y., Hao, S., Andersen-Nissen, E., Mauck, W. M., Zheng, S., Butler, A., Lee, M. J., Wilk, A. J., Darby, C., Zagar, M., Hoffman, P., Stoeckius, M., Papalexi, E., Mimitou, E. P., Jain, J., Srivastava, A., Stuart, T., Fleming, L. B., Yeung, B., … Satija, R. (2020). Integrated analysis of multimodal single-cell data. Cell, 184, 3573.

Heumos, L., & Theis, F. (2021). Ehrapy: Exploratory analysis of electronic health records. https://github.com/theislab/ehrapy

Hoyer, S., & Hamman, J. (2017). Xarray: ND labeled arrays and datasets in python. Journal of Open Research Software, 5(1), 10.

Huber, W., Carey, V., Gentleman, R., Anders, S., Carlson, M., Carvalho, B., Bravo, H., Davis, S., Gatto, L., Girke, T., Gottardo, R., Hahne, F., Hansen, K., Irizarry, R., Lawrence, M., Love, M., MacDonald, J., Obenchain, V., Oleś, A., … Morgan, M. (2015). Orchestrating high-throughput genomic analysis with bioconductor. Nature Methods, 12(2), 115–121.

Mangiola, S. (2021). tidySummarizedExperiment: Brings SummarizedExperiment to the tidy-verse. Bioconductor. https://doi.org/10.18129/B9.bioc.tidySummarizedExperiment

McInnes, L., Healy, J., & Melville, J. (2020). UMAP: Uniform manifold approximation and projection for dimension reduction. arXiv, 1802.03426.

McKinney, W. (2010). Data structures for statistical computing in python: Vols. Proceedings of the 9th Python in Science Conference 445 (pp. 51–56). Austin, TX.

Megill, C., Martin, B., Weaver, C., Bell, S., Prins, L., Badajoz, S., McCandless, B., Pisco, A. O., Kinsella, M., Gri in, F., Kiggins, J., Haliburton, G., Mani, A., Weiden, M., Dunitz, M., Lombardo, M., Huang, T., Smith, T., Chambers, S., … Carr, A. (2021). Cellxgene: A performant, scalable exploration platform for high dimensional sparse matrices. bioRxiv. https://doi.org/10.1101/2021.04.05.438318

Miles, A., Kirkham, J., Durant, M., Bourbeau, J., Onalan, T., Hamman, J., Patel, Z., shikharsg Rocklin, M., dussin, raphael, Schut, V., Andrade, E.S. de, Abernathey, R., Noyes, C., sbalmer, bot, pyup.io, Tran, T., Saalfeld, S., Swaney, J., … Banihirwe, A. (2020). Zarr. Zenodo. https://doi.org/10.5281/zenodo.3773450

Palla, G., Spitzer, H., Klein, M., Fischer, D., Schaar, A. C., Kuemmerle, L. B., Rybakov, S., Ibarra, I. L., Holmberg, O., Virshup, I., Lotfollahi, M., Richter, S., & Theis, F. J. (2021). Squidpy: A scalable framework for spatial single cell analysis. bioRxiv. https://doi.org/10.1101/2021.02.19.431994

Papadopoulos, S., Datta, K., Madden, S., & Mattson, T. (2016). The tiledb array data storage manager. Proceedings of the VLDB Endowment, 10(4), 349–360.

Paszke, A., Gross, S., Massa, F., Lerer, A., Bradbury, J., Chanan, G., Killeen, T., Lin, Z., Gimelshein, N., Antiga, L., Desmaison, A., Kopf, A., Yang, E., DeVito, Z., Raison, M., Tejani, A., Chilamkurthy, S., Steiner, B., Fang, L., … Chintala, S. (2019). PyTorch: An imperative style, high-performance deep learning library. In Advances in neural information processing systems 32 (pp. 8024–8035).

Pedregosa, F., Varoquaux, G., Gramfort, A., Michel, V., Thirion, B., Grisel, O., Blondel, M., Prettenhofer, P., Weiss, R., Dubourg, V., Vanderplas, J., Passos, A., Cournapeau, D., Brucher, M., Perrot, M., & Duchesnay, E. (2011). Scikit-learn: Machine learning in Python. Journal of Machine Learning Research, 12, 2825–2830.

Ramos, M., Schiffer, L., Re, A., Azhar, R., Basunia, A., Rodriguez, C., Chan, T., Chapman, P., Davis, S. R., Gomez-Cabrero, D., Culhane, A. C., Haibe-Kains, B., Hansen, K. D., Kodali, H., Louis, M. S., Mer, A. S., Riester, M., Morgan, M., Carey, V., & Waldron, L. (2017). Software for the integration of multiomics experiments in bioconductor. Cancer Research, 77 (21), e39–e42.

Regev, A., Teichmann, S. A., Lander, E. S., Amit, I., Benoist, C., Birney, E., Bodenmiller, B., Campbell, P., Carninci, P., Clatworthy, M., & others. (2017). Science forum: The human cell atlas. Elife, 6, e27041.

Wickham, H. (2014). Tidy data. Journal of Statistical Software, 59 (10), 1–23.

Wolf, F. A., Angerer, P., & Theis, F. J. (2018). SCANPY: Large-scale single-cell gene expression data analysis. Genome Biology, 19 (1), 15.

Zappia, L., & Theis, F. J. (2021). Over 1000 tools reveal trends in the single-cell RNA-seq analysis landscape. Genome Biology, 22(1), 1–18.

